# Heavy Metal Pollution From a Major Earthquake and Tsunami in Chile Is Associated With Geographic Divergence of Clinical Isolates of Methicillin-Resistant *Staphylococcus aureus* in Latin America

**DOI:** 10.1101/2023.05.18.541300

**Authors:** Jose RW Martínez, Manuel Alcalde-Rico, Estefanía Jara-Videla, Rafael Rios, Ahmed M. Moustafa, Blake Hanson, Lina Rivas, Lina P. Carvajal, Sandra Rincon, Lorena Diaz, Jinnethe Reyes, Ana Quesille-Villalobos, Roberto Riquelme-Neira, Eduardo A. Undurraga, Jorge Olivares-Pacheco, Patricia García, Rafael Araos, Paul J. Planet, César A. Arias, Jose M. Munita

## Abstract

Methicillin-resistant Staphylococcus aureus (MRSA) is a priority pathogen listed by the World Health Organization. The global spread of MRSA is characterized by successive waves of epidemic clones that predominate in specific geographical regions. The acquisition of genes encoding resistance to heavy-metals is thought to be a key feature in the divergence and geographical spread of MRSA. Increasing evidence suggests that extreme natural events, such as earthquakes and tsunamis, could release heavy-metals into the environment. However, the impact of environmental exposition to heavy-metals on the divergence and spread of MRSA clones has been insufficiently explored. We assess the association between a major earthquake and tsunami in an industrialized port in southern Chile and MRSA clone divergence in Latin America. We performed a phylogenomic reconstruction of 113 MRSA clinical isolates from seven Latin American healthcare centers, including 25 isolates collected in a geographic area affected by an earthquake and tsunami that led to high levels of heavy-metal environmental contamination. We found a divergence event strongly associated with the presence of a plasmid harboring heavy-metal resistance genes in the isolates obtained in the area where the earthquake and tsunami occurred. Moreover, clinical isolates carrying this plasmid showed increased tolerance to mercury, arsenic, and cadmium. We also observed a physiological burden in the plasmid-carrying isolates in absence of heavy-metals. Our results are the first evidence that suggests that heavy-metal contamination, in the aftermath of an environmental disaster, appears to be a key evolutionary event for the spread and dissemination of MRSA in Latin America.

## Introduction

Methicillin-resistant *Staphylococcus aureus* (MRSA) infections are a major global public health problem, with the World Health Organization considering the development of new therapeutic alternatives against MRSA a top priority (WHO, 2017; Murray et al. 2022). MRSA has distinctive epidemiologic patterns with specific genetic lineages restricted to particular geographical areas.(Arias et al. 2017; Challagundla et al. 2018). The widespread dissemination of MRSA over time is often driven by “waves” of clonal replacements, where novel lineages replace predominant regional clones (Planet et al. 2017). While the underlying factors driving clonal replacement of MRSA remain unclear, this phenomenon has been widely reported in different parts of the world, including Latin America. One of the most successful MRSA lineages in Latin America has been a healthcare-associated, ST5-SCC*mec*I lineage, which was first described in 1998 in Chile and Argentina (designated Chilean-Cordobes clone, [ChC])(Medina et al. 2013; Martínez et al. 2019). By the end of the 2000s, however, this lineage was almost completely replaced by a community-acquired MRSA (CA-MRSA) clone identified as USA300-Latin American variant (USA300-LV) in Colombia and Ecuador. In contrast, the ChC clone has remained largely dominant in countries like Chile and Peru, located on the South Pacific coast of Latin America (Aires de Sousa et al. 2001; Reyes et al. 2009; Arias et al. 2017).

USA300-LV belonged to an ST8-SCC*mec*IVc lineage that was closely related to the CA-MRSA USA300 clone (ST8-SCC*mec*IVa) responsible for a major epidemic of CA-MRSA infections in the United States in the late 1980s and 1990s (Planet et al. 2015). Interestingly, the evolutionary divergence observed between these lineages was associated with the independent acquisition of two horizontally-acquired genetic elements: the <u>A</u>rginine <u>C</u>atabolic <u>M</u>obile <u>E</u>lement (ACME) in the North American USA300 clone, and the <u>Co</u>pper (Cu) and <u>Mer</u>cury (Hg) resistance mobile element (COMER) in South American lineage (Planet et al. 2015). Of note, apart from the arginine metabolism machinery, ACME also harbored *copX*(B), a Cu resistance gene also observed in other successful MRSA clones (Saenkham-Huntsinger et al. 2021). Furthermore, previous studies have suggested a possible evolutionary advantage of acquiring heavy metal resistance traits in the emergence of new MRSA lineages (Kernberger-Fischer et al. 2018; Zapotoczna et al. 2018). These observations suggest that acquiring mobile genetic elements harboring heavy-metal resistance genes (HMRGs) might play a key role in the emergence and dissemination of successful MRSA clones. However, the possible underlying environmental causes driving the appearance and dissemination of MRSA lineages remain unclear.

Studies involving non-pathogenic bacteria suggest environmental contamination with heavy metals promotes horizontal gene transfer of antimicrobial resistance genes and selects for organisms harboring plasmids that carry heavy metal resistance traits, which are frequently co-transferred with other antimicrobial resistance determinants (Xu et al. 2017; Rodgers et al. 2018; Zhang et al. 2018). In addition, extreme natural events such as volcano eruptions, heavy rainfalls, earthquakes, and tsunamis release high amounts of heavy metals into the environment (Shruti et al. 2018; Brizuela et al. 2019; Ji et al. 2021; Ota et al. 2021). However, the potential role of such events as drivers of the evolution of clinically-relevant antimicrobial-resistant pathogens remains unclear.

On February 27, 2010, the sixth-largest earthquake ever recorded (Mw 8.8) occurred off the coast of central Chile. The earthquake, which was also felt in some parts of Argentina and Peru, triggered a subsequent tsunami and landslides that affected several coastal towns and cities. The disaster resulted in about 0.5 million homes damaged, thousands of people injured, 525 deaths, and 23 people missing. It also had a major environmental impact. The tsunami severely damaged the industrialized port of Talcahuano, Concepción Bay, which includes an oil refinery, steel, and cement production, petrochemical industries, coal power stations, and military and civilian shipyards. Data suggest that the disaster resulted in a higher-than-average concentration of heavy metals in marine sediments, urban soils, and marine fauna (Luz María Fariña; Cristián Opaso; Paulina Vera 2012; Tume et al. 2018; Tapia et al. 2019).

In this study, we aimed to explore the possible role of the 2010 earthquake and tsunami, and the release and resuspension of heavy metals into the environment in the industrialized coastline of Concepción, as a driving force for the selection and evolution of MRSA genomic lineages circulating in Chile. We performed a detailed phylogenomic reconstruction of 113 ChC clone MRSA clinical isolates recovered from bloodstream infections,(Arias et al. 2017) obtained from seven healthcare centers in six countries in Latin America, including one in Concepción, Chile. We used hybrid assemblies combining short- and long-read sequencing to evaluate the potential impact of mobile genetic elements carrying horizontally transferable Heavy Metal Resistance Genes (HMRGs) in the evolution of the MRSA ST5-SCC*mec*I ChC clone.

## Results

### Characteristics of the ChC MRSA strains collection

Our study included 113 ChC MRSA bacteremia isolates recovered from seven hospitals in Lima, Perú (n=37, 32.7%); Santiago, Chile (n=28, 24.8%); Concepción, Chile (n=25, 22.1%); Caracas, Venezuela (n=13, 11.5%); Sao Paulo, Brazil (n=5, 4.4%); Bogotá, Colombia (n=3, 2.7%); and Buenos Aires, Argentina (n=2, 1.8%). As typically described for the ChC MRSA clone, isolates exhibited high rates of resistance to ciprofloxacin, gentamicin, erythromycin, and clindamycin, along with susceptibility to tetracyclines, cotrimoxazole, rifampicin, vancomycin, and linezolid (Table S1). A total of 27% of the isolates were susceptible to ceftaroline, while the remaining 73% exhibited a minimal inhibitory concentration in the susceptible dose-dependent range, as per CLSI breakpoints (Table S1), consistent with a previous report (Khan et al. 2019). We observed no statistically significant differences in antimicrobial susceptibility across geographical locations (Table S1).

### WGS analyses and phylogeographical relatedness of ChC MRSA in LA

The *in-silico* sequence type (ST) determination revealed that all the isolates belonged to clonal complex 5 and identified 112 out of the 113 isolates (99%) as ST5; the remaining isolate was classified as ST105. Out of the 112 ST5 strains, 109 (97%) were considered ChC clones (carried *mecA* on SCC*mec*I); the remaining three isolates harbored a non-classical ChC clone SCC*mec*IV cassette. The ST105 isolate carried *mecA* in SCC*mec*II.

Our core genome-based phylogenomic reconstruction revealed that the 109 genomes belonging to the ST5-SCC*mec*I (ChC) clone grouped into three well-defined clades that followed a marked geographic pattern (Fig. 1). Clade I (ChC-I, n=9) predominantly consisted of MRSA isolates from Chile (n=8), with one from Peru. Clade II (ChC-II, n=28) only contained isolates recovered from patients in Chile. Finally, clade III (ChC-III, n=72), the largest and most diverse, was further split into three sub-clades: i) ChC-IIIa (n=21) including 16 isolates from Chile and all five strains from Brazil; ii) ChC-IIIb (n=16) grouped all isolates from Venezuela (n=10), Colombia (n=3) and Argentina (n=2), along with one from Peru; and iii) ChC-IIIc (n=35) only included isolates recovered from Peru (Fig. 1).

**Figure 1.**
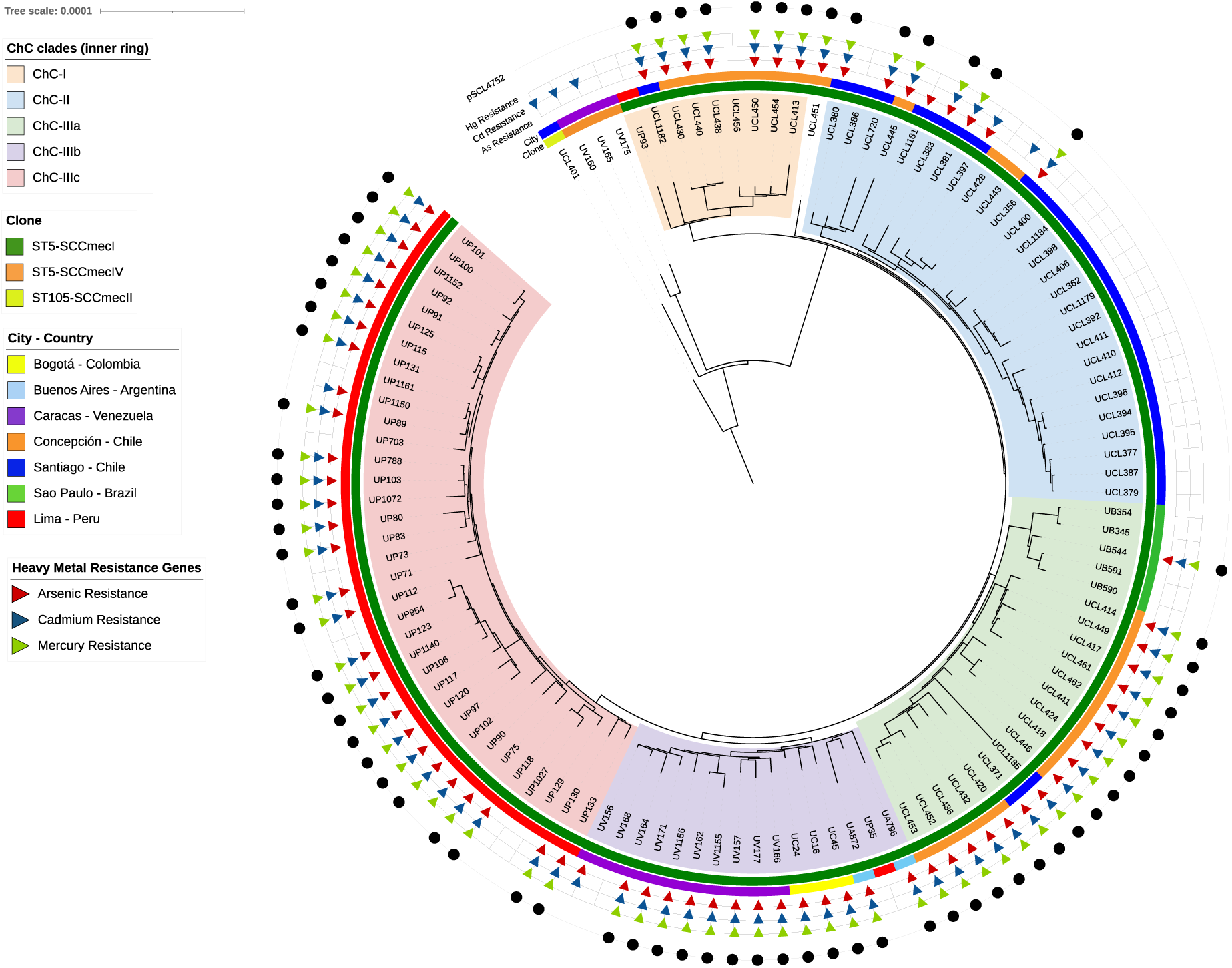
Core genome based phylogenomic reconstruction of the 113 Latin American ChC clone MRSA genomes. The phylogenomic reconstruction was rooted at the midpoint of genomic distances. The most important clades are represented by colors within the reconstruction. The inner colored ring indicates the ST-SCC*mec*. The outer colored ring indicates the city of origin of the isolates. The outer triangles show the presence of the heavy-metal resistance genes: red, arsenic resistance genes (*arsB* and/or *arsC*); blue, cadmium (*cadA*, *cadC*, and/or *cadD*); and green, mercury (*merA*, *merB, merT,* and/or *merR*).

### High prevalence of pSCL4752 plasmid-encoded HMRGs in Latin American ST5-SCCmecI MRSA

To evaluate the potential role of HMRGs in the divergence of ST5-SCC*mec*I MRSA, we performed an *in silico* search of horizontally-acquired HMRGs involved in the processing of heavy metals. A total of 80/113 (71%) isolates harbored at least one set of acquired HMRGs associated with resistance to As (*arsBC*), Cd (*cadACD*), or Hg (*merABTR*). Among them, 72 out of 80 (90%) co-carried all the resistance determinants for As, Cd, and Hg (Fig. 1). Of the remaining eight isolates, three carried genes encoding resistance to As and Cd, four only carried As resistance genes, and one harbored cadmium resistance genes alone. Interestingly, all 113 isolates lacked *copX(B),* an acquired determinant involved in Cu resistance previously found in other MRSA clones.

Since co-detection of horizontally acquired HMRGs was frequently observed among our isolates (Fig. 1), we sought to determine the genomic context of these HMRGs by performing a hybrid assembly (short-read and LRS) of a representative strain (SCL 4752) harboring As, Cd, and Hg resistance traits. Our results generated a complete genome of 3.052.503 bp with a GC content of 32.9%, composed of two contigs, including the chromosome and a plasmid that we designated pSCL4752. This plasmid consisted of a total of 36,660 bp with a GC content of 32.5% (Fig. 2), and shared extensive identity (99.9%) with a rep20_3_rep(pTW20)/rep21_20_p020(pLGA251) plasmid designated pCM05. Of note, pCM05 had been previously identified in a linezolid-resistant ST5-SCC*mec*I MRSA strain isolated in Colombia (NC_013323.1)(Arias et al. 2008). pSCL4752 was predicted to encode a total of 41 CDSs, including all horizontally-acquired HMRGs described above (*arsBC, cadACD, merABTR*), and a copy of the *blaIRZ* operon, which encodes the expression of the staphylococcal penicillinase, BlaZ (Fig. 2). pSCL4752 also contained two duplicated invertases (*bin3* and *hin*), five transposases (*IS431L*, *IS431R*, *ISSau6*, *ISBli29*, IS481), and a plasmid replication initiator protein (*repA*), along with three replication proteins and several hypothetical proteins (Fig. 2). *merABTR*, *resA*, *garB*, and one hypothetical protein were flanked by two IS26 family transposases (*IS431L* and *IS431R*), potentially suggesting the presence of a mobile mercury resistance transposon.

**Figure 2.**
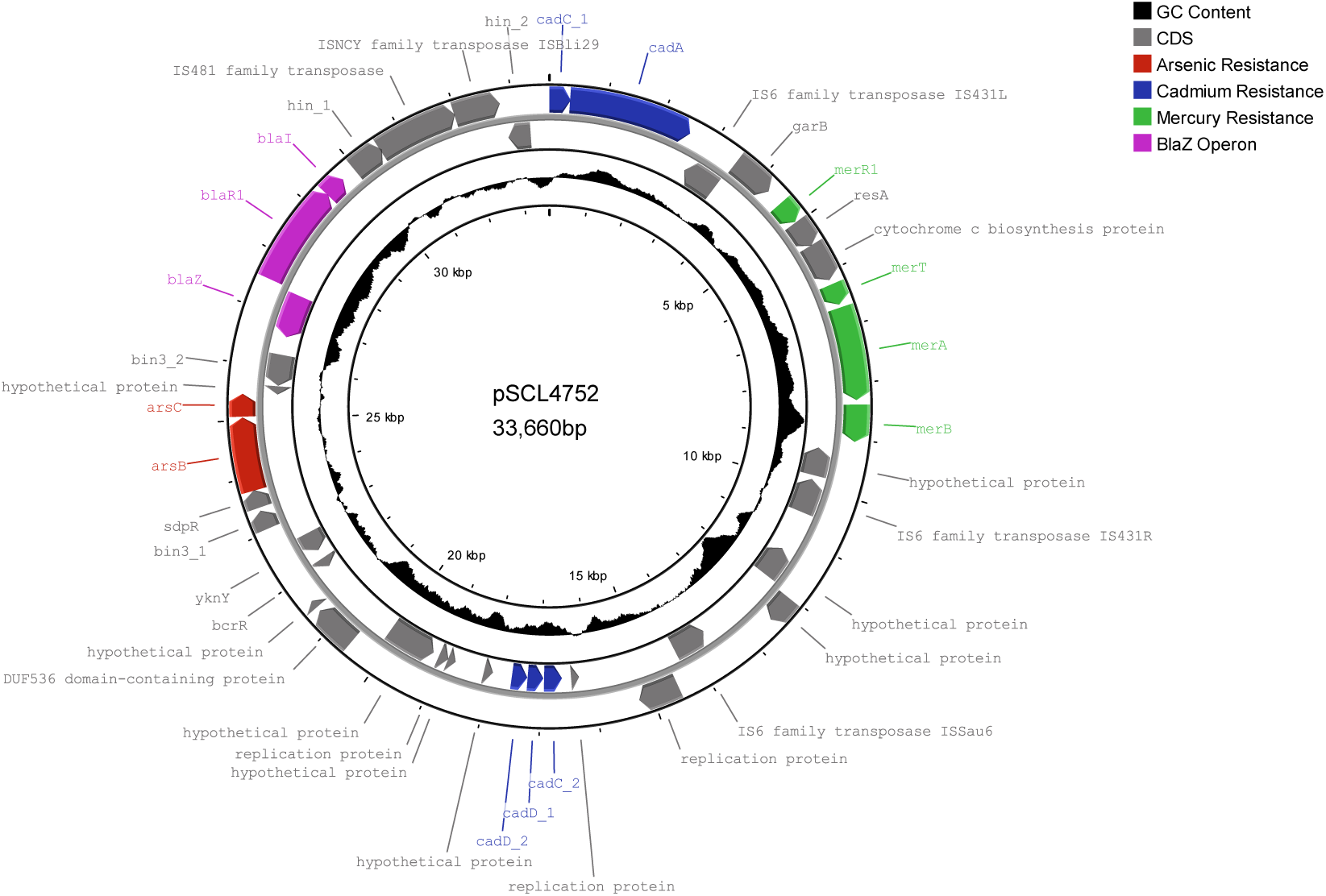
Schematic representation of plasmid pSCL4752. **The** Circularized pSCL4752 plasmid includes the annotations of the coding sequences (gray arrows). Heavy-metal resistance genes for arsenic, cadmium, and mercury are shown in colored arrows (red, blue, and green, respectively). The *blaZ* operon is depicted in purple.

### Geographical divergence of the ST5-SCCmecI MRSA clone in Chile is associated with the presence of pSCL4752

A total of 71 out of the 113 (63%) isolates harbored the pSCL4752 plasmid. Noteworthy, the frequency of this plasmid in clades ChC-I (77%) and ChC-III (81%) strongly varied compared to clade ChC-II (35%) (Fig 1). Interestingly, clade ChC-II was mainly composed of isolates recovered from Santiago (central Chile). In contrast, the isolates grouped in clades ChC-I and ChC-III were obtained from Concepción (southern Chile). Thus, we aimed to study the possible role of pSCL4752 as a driver of MRSA evolutionary divergence. Since the major divergence was observed in MRSA isolates from Chile, we focused the analysis on the genomes of the 53 Chilean MRSA isolates recovered from Santiago and Concepción.

An in-depth Bayesian molecular clock analysis using the 53 Chilean genomes estimated the most recent common ancestor in 2008 (95% high posterior density interval [HPD] 2007.03-2008.77) (Fig. 3). The molecular clock revealed a major divergence event in March 2010 (following the February 27 earthquake), which was quickly followed by two secondary divergence events occurring in parallel between September and November of 2010. As shown in Fig. 3, these events grouped isolates into four clades highly associated with the city of origin (Santiago and Concepción). This geographical divergence was also linked to the presence of pSCL4752 (Fisher’s exact test p<0.0001) (Fig. 3). Indeed, the prevalence of carriage of pSCL4752 was 88% for isolates recovered from Concepción and only 29% for Santiago. These results suggest that the environmental heavy metal pollution associated with the 2010 earthquake and subsequent tsunami was a major driver of the geographic divergence observed in Chilean ST5-SCC*mec*I MRSA.

**Figure 3.**
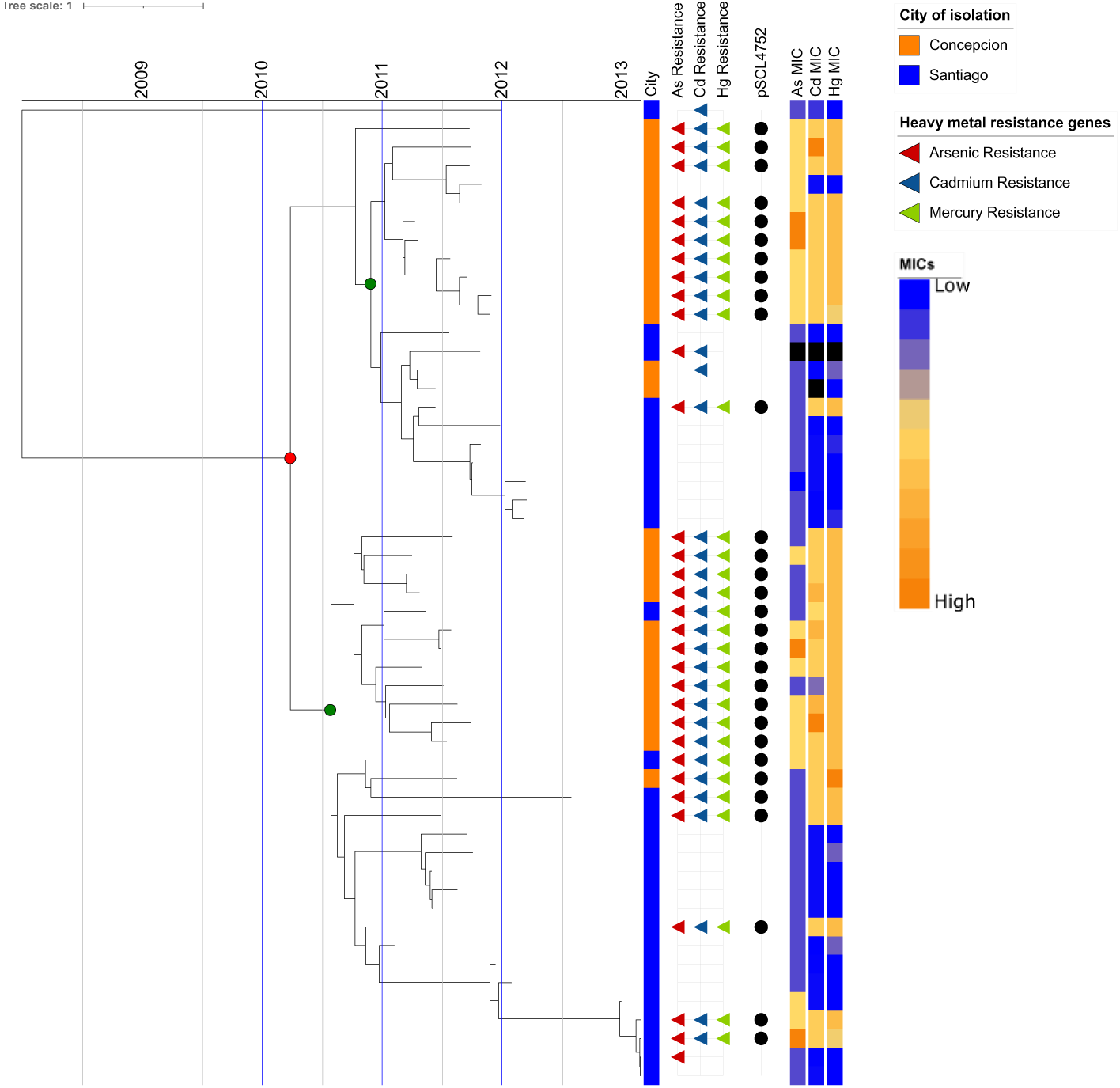
Bayesian phylogenomic reconstruction of the 53 ST5-SCC*mec*I isolates collected between 2010-2013 in Chile. The tips of each branch of the tree correspond to the isolation date and the time scale is displayed at the top of the tree. The colored circles in the tree represent the main divergence events indicating the expansion of the ChC clone (red) and the parallel divergence (green). The colored band shows the city of origin of the isolation. The outer triangles show the presence of the heavy-metal resistance genes: red, arsenic resistance genes (*arsB* and/or *arsC*); blue, cadmium (*cadA*, *cadC*, and/or *cadD*); and green, mercury (*merA*, *merB, merT,* and/or *merR*). The presence of plasmid pSCL4752 is indicated by black circles. The heatmap shows the minimum inhibitory concentration to arsenic (12.5-400 µM NaAsO2), cadmium (6.25-3200 µM CdSO4) and mercury (1.5-100 µM HgSO4) for each strain tested.

### Isolates harboring the pSCL4752 plasmid exhibited increased resistance to heavy metals

To assess the functionality of the HMRGs contained in pSCL4752, we performed susceptibility testing for As, Cd, Hg, and Cu by broth microdilution in the 53 Chilean isolates (Figs. 3 and 4). Overall, plasmid-harboring strains exhibited significantly higher MICs to Hg, Cd, and As (p<0.0001). Indeed, the MIC_50/90_ for Hg, Cd, and As in isolates harboring pSCL4752 were 25/25µM, 800/1600µM, and 100/200µM, respectively, and 1.5/6.25µM, 25/25µM, and 50/100µM, in those lacking the plasmid (Fig. 4). In concordance with the absence of Cu resistance genes on the plasmid, the MIC_50/90_ values to Cu did not vary between strains with or without pSCL4752 (Fig. 4). Hence, our results support the notion that MRSA isolates harboring pSCL4752 could be positively selected in environments with high concentrations of heavy metals, such as observed after the 2010 tsunami in the coast of Concepción in southern Chile.

**Figure 4.**
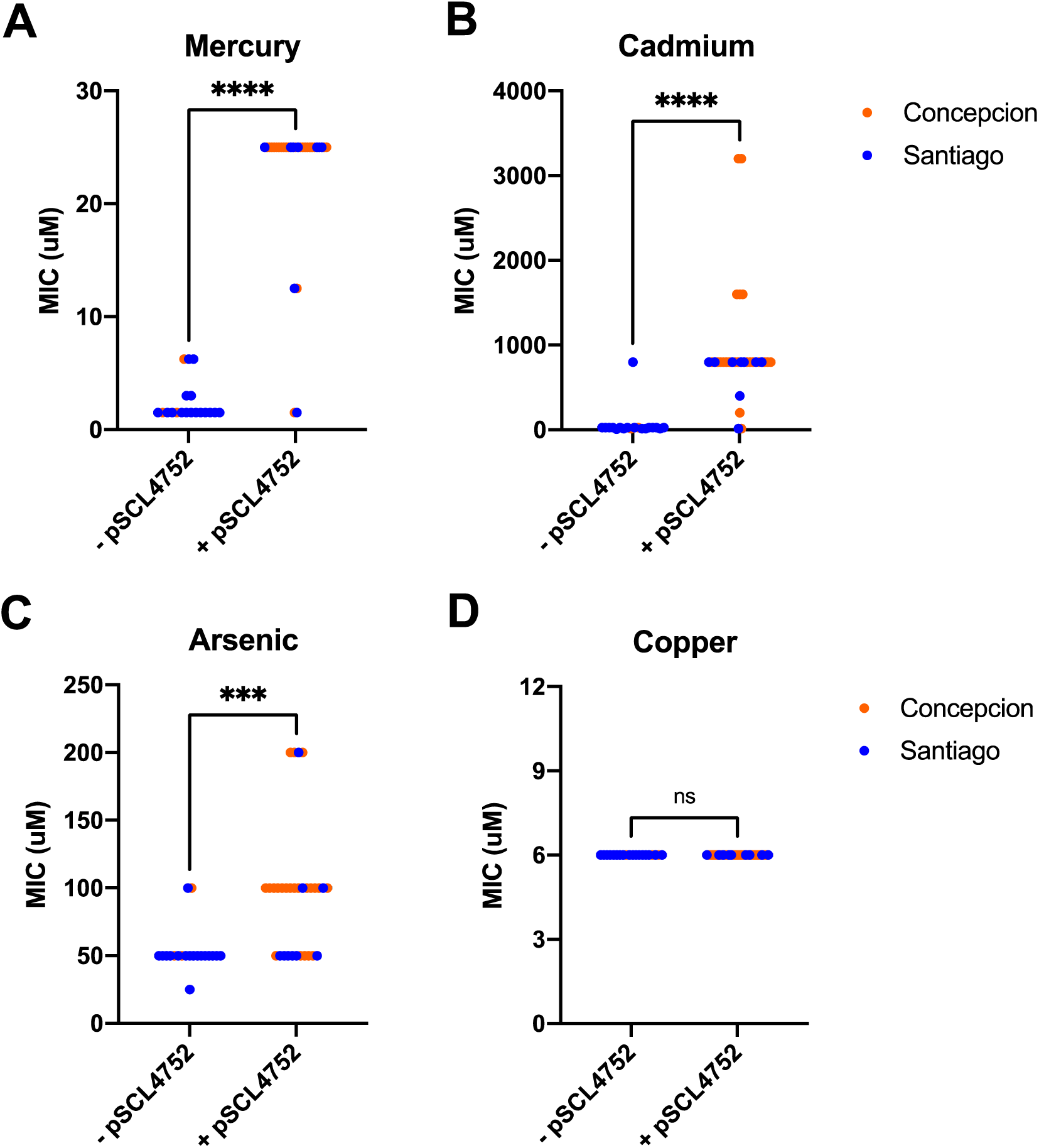
Phenotypical effect of the presence of the pSCL4752 plasmid in Chilean clinical isolates. Broth microdilution MICs of the 53 Chilean clinical isolates to mercury (A), cadmium (B), arsenic (C), and copper (D). The MIC value was determined as the minimal concentration that inhibits bacterial growth. Statistical analysis was performed with the non-parametric Mann-Whitney test. *p <0.05, ***p < 0.001, ****p <0.0001, ns = non-significant.

### The presence of pSCL4752 carries a fitness cost in the absence of heavy metals

To determine the role of pSCL4752 in resistance to heavy metals and to evaluate a possible fitness cost associated with its carriage, the plasmid was “cured” in one representative strain from each of the four clades established by the molecular clock analysis (Fig. 3). The loss of the plasmid was observed in all the isolates after two days of growth in trypticase soy broth medium, which was confirmed by polymerase chain reaction. All isogenic strains in which pSCL4752 was cured presented a statistically significant reduction in the minimal inhibitory concentrations of Hg (p=0.001), Cd (p=0.0005), and As (p=0.0313), as compared to their plasmid-harboring counterparts (Figs. 5A-C). In addition, plasmid-cured strains grew faster (average doubling time 42 min±5.7 vs. 59 min±3.8, respectively; p=0.0005) and reached a higher OD_600_ (1.638 ±0.1; p<0.05) than their isogenic parental strains harboring the plasmid (Fig. 5, lower panel). These results suggest that the presence of pSCL4752 confers an evolutionary advantage to ChC isolates in the presence of sub-inhibitory concentrations of heavy metals but introduces a fitness cost when this selection pressure is removed.

**Figure 5.**
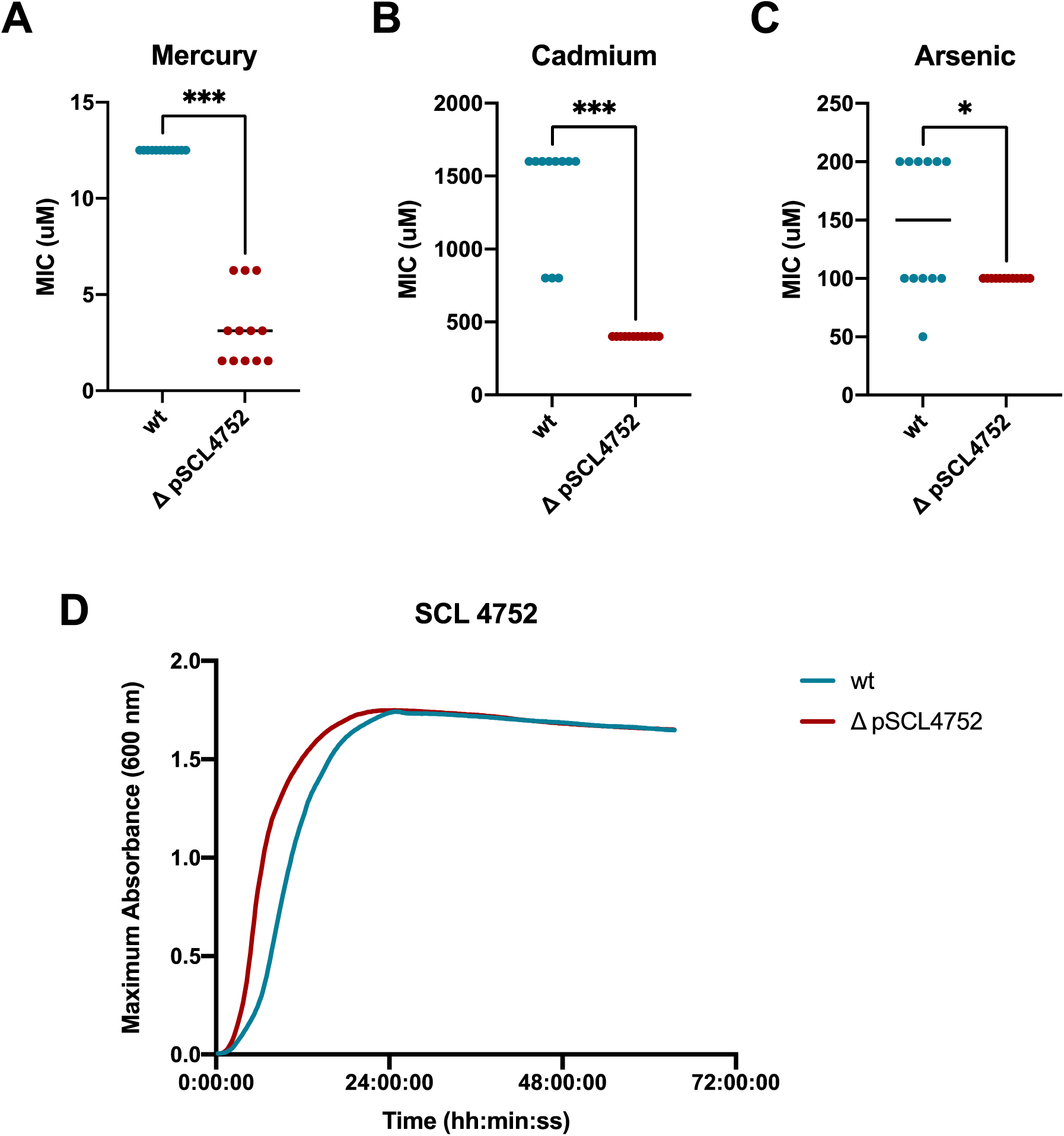
Phenotypical analysis of four pSCL4752 cured strains. MIC determination by BMD method to mercury (A), cadmium (B), arsenic (C) in four MRSA isogenic clone strains carrying the plasmid (wt), and plasmid cured (ΔpSCL4752). The MIC value were determined as the minimal concentration that inhibits bacterial growth. D. Growth curve of representative plasmid-cured strain. The color of the curves represents the plasmid curing treatment, being dark red for treated and green for non-treated. The X axis shows the time and the Y axis the OD_600_. All the curves were performed in technical triplicates from at least two independent experiments. Statistical analysis was performed with the non-parametric Wilcoxon matched pairs signed rank test. *p <0.05, ***p < 0.001, ****p <0.0001, ns = non-significant.

## Discussion

Increasing evidence suggests an association between the acquisition of heavy metal resistance determinants and the rise of antimicrobial-resistant pathogens (Xu et al. 2017; Zhang et al. 2018; Biswas et al. 2021). Indeed, the chromosomal acquisition of horizontally-transferred HMRGs has been linked to the evolutionary divergence of North and South American epidemics of USA300-LV and USA300, two major CA-MRSA clones (Planet et al. 2015). However, data on the potential role of plasmids harboring HMRGs as drivers of the evolution and spread of clinically relevant MRSA lineages are scant. Herein, we describe a major evolutionary divergence event in clinical isolates of the ST5-SCC*mec*I ChC clone MRSA that was associated with the presence of a plasmid (pSCL4752), harboring heavy metal resistance determinants. This divergence, estimated to have occurred in 2010, followed a distinct geographical distribution, clustering isolates recovered from Santiago and Concepción, two Chilean cities. Additionally, a unique phylogeographic analysis of the ST5-SCC*mec*I MRSA clone in Latin America revealed a distinct geographic clustering highly associated with the country of bacterial isolation.

Our results align with previous studies describing the emergence of new MRSA lineages associated with possible evolutionary advantages of heavy metal resistance traits (Kernberger-Fischer et al. 2018; Zapotoczna et al. 2018). Environmental contamination with heavy metals has been associated with horizontal gene transfer and the selection of non-pathogenic organisms harboring plasmids that carry heavy metal resistance traits (Xu et al. 2017; Zhang et al. 2018). The divergence observed could be partly driven by environmental selective pressure. Indeed, historical records report high levels of heavy metal pollution in urban soils and marine sediments in Concepción (Barrios-Guerra 2004; Tume et al. 2008; Luz María Fariña; Cristián Opaso; Paulina Vera 2012; Tume et al. 2018). Research has shown that tsunamis and other major catastrophic events release and resuspend heavy metals from marine sediments and land-based pollutants (Brizuela et al. 2019; Ota et al. 2021). A previous study found a significant increase in heavy metals observed in mollusks collected off the coast of Concepción following the 2010 tsunami (Tapia et al. 2019). Our molecular clock analyses estimated that the initial divergence event leading to the selection of HMRGs-harboring plasmid pSCL4752 occurred between March and September 2010. Altogether, these data suggest that the increase in environmental heavy metals released by the 2010 earthquake and subsequent tsunami contributed to the selective pressure driving the divergence events observed in the ST5-SCC*mec*I MRSA clone.

Phenotypically, isolates containing pSCL4752 exhibited higher tolerance to As, Cd, and Hg, suggesting an active role of the HMRGs harbored on the plasmid. However, in the absence of heavy metals, clinical isolates containing pSCL4752 exhibited slower growth than those not carrying the plasmid, suggesting that pSCL4752 could provide an evolutionary advantage in environments containing heavy metals, increasing the survivability of MRSA. On the other hand, maintenance of pSCL4752 in the absence of heavy metals resulted in a fitness burden. Interestingly, previous data have shown some MRSA clones have maintained mobile genetic elements containing HMRGs despite a fitness cost, due to an adaptative advantage beyond heavy metal resistance. Indeed, a horizontally transferred copper-resistant locus provided increased survival in macrophages and was associated with co-carriage of crucial antimicrobial resistance determinants in USA300 (Zapotoczna et al. 2018; Rosario-Cruz et al. 2019). We detected a *blaIRZ* operon present in pSCL4725, suggesting it may be related to the selection of the plasmid. However, we found the *blaIRZ* operon in 57% (n=24) of the isolates not carrying the pSCL4725 plasmid, suggesting that the plasmid was most likely selected by heavy metals and not by a potential advantage provided by this antimicrobial resistance operon. Furthermore, our genomic analyses revealed that Hg resistance is likely transposable since Hg resistance genes were contained within a transposon-like structure flanked by two IS26 family transposases (*IS431L* and *IS431R*). This element has been found in the chromosome linked to the SCC*mec* element in other MRSA lineages, including the COMER element (Planet et al. 2015). This, further suggest that mobile Hg resistance determinants might play a major role in the selection of successful MRSA lineages.

Our core genome-based phylogeographic analyses of isolates belonging to the ST5-SCC*mec*I ChC clone MRSA revealed a substantial genomic heterogeneity strongly associated with the city of origin. These results align with previous data suggesting an inherently higher geographical diversity in MRSA isolates belonging to clonal complex 5 (which includes ST5-SCC*mec*I) as compared to other MRSA lineages (Challagundla et al. 2018). Geographic genomic heterogeneity has also been observed in other MRSA lineages such as ST105 and ST239, both of which underwent marked divergence within different regions of Brazil (Botelho et al. 2019; Viana et al. 2021). The divergence events that generated the North and South American USA300 clones subsequently led to further rapid clonal expansion across different geographic regions (Reyes et al. 2009). Furthermore, the appearance of two predominant variants of the USA300 clone in an outbreak in New York suggested that MRSA clones may undergo genomic divergences even within the same geographical area and genetic lineage (Copin et al. 2019).

This study has some limitations. First, Chile was the only country where isolates were collected from two different cities. Therefore, we cannot discard the possibility that similar divergence events associated with the loss of pSCL4752 occurred in other regions of Latin America. A larger sample size with a more geographical and temporal representation of isolates would improve our understanding of the evolutionary history of the ST5-SCC*mec*I MRSA lineage and the impact of horizontally-acquired HMRGs, and the pSCL4752 plasmid in particular, on the evolution of this clone in Latin America. Second, there is no record of heavy metal concentrations in the clinical settings in Santiago or Concepción in the aftermath of the 2010 earthquake. Therefore, we cannot compare the minimum inhibitory concentrations obtained in this study with real-world conditions.

In conclusion, we used genomic data from clinical isolates of the ST5-SCC*mec*I ChC MRSA clone to describe a major evolutionary divergence event associated with plasmid-harbored heavy metal resistance genes. We observed that the divergence follows a spatiotemporal pattern probably associated with heavy metal pollution associated with an extreme natural event, the 2010 earthquake, and tsunami. Indeed, we found suggestive evidence of a possible link between the release of higher quantities of heavy metals in the aftermath of an environmental disaster and the divergent evolution of the ChC MRSA in the region. Improving our understanding of how chronic exposure and adaptation to environmental pollution associated with extreme events could affect the emergence of antimicrobial resistance determinants is critical for avoiding a potential future health crisis. Our results highlight the urgent need for additional research on environmental risk factors associated with the emergence of antimicrobial resistance.

## Materials and Methods

### Strain collection, antibiotic and heavy metal susceptibility testing

We studied a collection of 113 MRSA isolates recovered from adult patients diagnosed with *S. aureus* bacteremia between January 2011 and July 2014 in Argentina, Brazil, Chile, Colombia, Peru, and Venezuela. Susceptibility testing of antimicrobials and heavy metals was performed using either agar dilution or broth microdilution method according to the 2019 Clinical and Laboratory Standards Institute (CLSI) (details in Appendix 1).

### Whole-genome sequencing (WGS) and Phylogenomic analysis

Methods for DNA extraction, WGS, and *in silico* characterization for the 113 isolates are described in Appendix 1. The genome of a representative isolate (SCL 4752) was sequenced by long-read sequencing (LRS) (MinION, Oxford Nanopore Technologies, Oxford, UK) using the SQK-LSK208 kit following the manufacturer’s instructions and a consensus hybrid assembly using both Illumina and LRS reads was obtained. The phylogenomic relationships were assessed with a maximum likelihood phylogenomic tree and a Bayesian molecular clock using a GTR substitution model (Appendix 1).

### Plasmid curing and Growth curves

The plasmid curing protocol consisted of consecutive 24 hours passages of cultures growing with shaking at 44°C in tubes containing fresh MH broth (May et al. 1964). Growth curves were measured in three replicates for 24 hours at 37°C (Details in Appendix 1).

## Acknowledgments

We gratefully acknowledge the Molecular Genetics and Antimicrobial Resistance Unit, from Universidad El Bosque, Bogota, Colombia, for the isolates included in this study. This research was partly funded by the research grants FONDECYT 1171805, and FONDECYT 1211947. The computational infrastructure was provided by FONDEQUIP EQM150093. CAA was supported by NIH/NIAID grants K24AI121296, R01AI134637, R01AI148342-01, and P01AI152999-01.

